# Cellular Lipids Regulate the Conformational Ensembles of the Disordered Intracellular Loop 3 in β2 Adrenergic Receptor

**DOI:** 10.1101/2023.11.28.569080

**Authors:** Elizaveta Mukhaleva, Tianyi Yang, Fredrik Sadler, Sivaraj Sivaramakrishnan, Ning Ma, Nagarajan Vaidehi

**Affiliations:** Irell and Manella Graduate School of Biological Sciences, Beckman Research Institute of the City of Hope, Duarte, CA, USA; Department of Computational and Quantitative Medicine, Beckman Research Institute of the City of Hope, Duarte, CA, USA; Biochemistry, Molecular Biology and Biophysics Graduate Program, University of Minnesota, Minneapolis, MN, USA; Department of Genetics, Cell Biology and Development, University of Minnesota, Minneapolis, MN, USA

## Abstract

The structurally disordered intracellular loops (ICLs) of G protein-coupled receptors (GPCRs) play a critical role in G protein coupling. In our previous work, we used a combination of FRET-based and computational methodologies to show that the third intracellular loop (ICL3) modulates the activity and G protein coupling selectivity in GPCRs. In the current study, we have uncovered the role of several lipid components in modulating the conformational ensemble of ICL3 of the β2-adrenergic receptor (β2AR). Our findings indicate that phosphatidylinositol 4,5-bisphosphate (PIP2) in the inner leaflet of the membrane bilayer acts as a stabilizing anchor for ICL3, opening the intracellular cavity to facilitate G protein coupling. This interaction between PIP2 and ICL3 causes tilting of β2AR within the cellular membrane. Notably, this tilting of the receptor is supported by ganglioside GM3 stabilizing the extracellular loops on the outer leaflet of the bilayer, thereby exerting an allosteric effect on the orthosteric ligand binding pocket. Our results underscore the significance of lipids in modulating GPCR activity, proposing an allosteric mechanism that occurs through the receptor’s orientation within the membrane.

## Introduction

The activity of G protein coupled receptors (GPCRs) are often affected by the structures and chemistry of the lipid molecules that surround them in the membrane^1^. Both the types of lipids and relative composition of lipids in the bilayer influence the conformational ensemble of the receptor. Many studies have shown that different lipids play an important role in GPCR function^2–9^ . NMR studies of β2-adrenergic receptor (β2AR) in detergent micelles compared to reconstituted high density lipoprotein environment showed that the basal activity and exchange rates between inactive and active conformations is higher in detergents and in phospholipid bilayer containing HDL particles compared to cell membranes ^2,10,11^. Although it is known that cellular lipids have a definitive effect on receptor activity, the mechanism(s) by which lipid chemistry affects the receptor activity remain nebulous.

The three intracellular loops and the carboxy terminus of GPCRs play a critical role in coupling to G proteins and β-arrestins. The extracellular loops are also involved in not only regulating the ligand binding but also in G protein coupling^12,13^. However, receptor loop regions and N- and C-termini are largely unstructured and often not resolved in the three-dimensional structures making it particularly difficult to study their role in GPCR function using structural techniques. Recent studies using biophysical techniques have shown that the intrinsically disordered intracellular loop 3 (ICL3) and the carboxy terminus of GPCRs regulate the receptor coupling selectivity to G proteins^14,15^. In our previous study^14^ on β2AR, we found that ICL3 spans a broad ensemble of conformational states, ranging from “closed” states that block the binding site of the G protein to “open” states that allow for the G protein to bind ^14^. Thus, ICL3 regulates the access to the G protein coupling site in the receptor. Through this mechanism, ICL3 acts as a secondary regulator of β2AR activation as well as plays an important role in the G protein coupling strength and selectivity. Although it is evident from this study that ICL3 regulates the activity and signaling specificity of β2AR, the role of the membrane lipids in stabilizing the conformation of ICL3 is not known. In this study we have used extensive molecular dynamics simulations (22 μs) in multi-lipid bilayer model that mimics the plasma membrane to analyze the effects of different types of lipids on the conformational ensemble of ICL3 and thereby their effects on β2AR activation.

## Results

### ICL3 acts as a secondary regulator β2AR of activity by forming a cap in the G protein binding site

As detailed in Table S1, we previously performed 22 µs of all-atom molecular dynamics (MD) simulations on β2AR with agonist including the full length ICL3 in various starting conformations in a multi-lipid bilayer comprising lipids consisting of POPC, DOPC, POPE, DOPE, POPS, DOPS, sphingomyelin (Sph), ganglioside (GM3), and cholesterol (CHOL) and phosphatidylinositol 4,5-bisphosphate (PIP2)^14,16^. We observed that ICL3 spanned a broad conformational ensemble in these simulations, with transitions between closed and open ensemble of conformations. The closed state of ICL3 is defined as a state where the C-terminus of the ICL3 blocks the entry of α-helix in the C-terminus of the G protein. To enrich the conformations near the transition of closed to open state of ICL3, we performed a swarm of MD simulations by initiating simulations from multiple population density maxima along the transition pathway. The total MD simulation trajectories sum up to 22 μs long. We sought to further analyze the role of the lipid components in stabilizing the different conformational states of ICL3 thereby regulating the activity of β2AR.

We performed Principal Component Analysis (PCA) using the backbone atom coordinates of β2AR and projected the MD snapshots on the top two scoring PCs(Fig. 1A). Each maxima on the PC landscape represents a potential representative state of the system, capturing key conformational states within the trajectory. Next, we clustered the MD frames using their first two PCs (see Methods) and obtained 12 distinct conformation clusters as depicted in Figs. S1A, S1B and S1C. By overlapping Fig 1A with Fig. S1C, it became apparent that 10 out of 12 clusters from Fig S1C corresponded with local maxima on the population density landscape in Fig 1A. Therefore, we focused our attention on those 10 clusters. We selected the ensemble of conformations around the 10 maxima for further analysis (see Methods).

**Figure 1.**
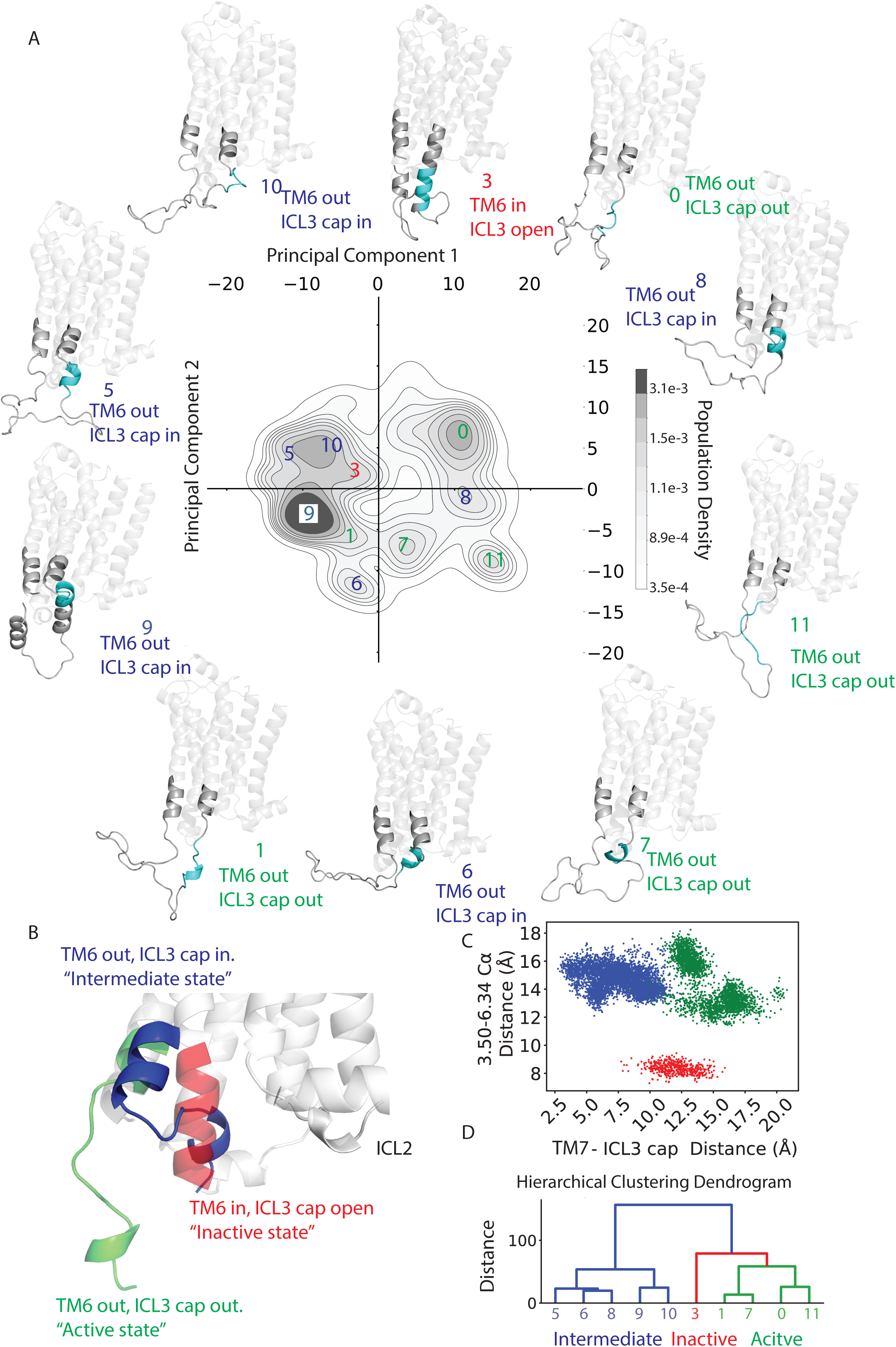
ICL3 acts as a secondary regulator of β2AR activity. A) Population density landscape showing the conformational ensembles of β2AR-ICL3. The population density landscape is shown in the principal component space (PC1 and PC2). The representative structures (colored grey) of the 10 distinct conformational clusters are shown. The ICL3 region is colored in dark grey, the ICL3 cap is colored in cyan. The conformational clusters of the active states are labeled in green; the intermediate states are labeled in blue, and the inactive states are labeled in red. **B**) Structure overlay to show the positions of TM6 and ICL3 in the inactive (red) intermediate (blue) and active (green). In the inactive state structure (red), TM6 is close to TM3 blocking the G protein binding site, and the ICL3 cap is open and making contacts with ICL2 and ICL1 residues. In the intermediate structure (blue), TM6 has moved outwards, ICL3 cap partially obstructs the G protein binding site. Both TM6 and ICL3 cap are out in the active state (green). Only the cap region of ICL3 is shown for clarity. **C**) Projection of all MD snapshots using the 3.50 and 6.34 distance and ICL3 cap-TM7 distance. **D**) The hierarchical clustering dendrogram of the ten clusters, using 3.50-6.34 distance and ICL3 cap-TM7 distance to separate the ten clusters into three major states.

The 10 conformational clusters are differentiable by two specific structural features of the receptor: the transmembrane helix (TM) 3-TM6 distance (3.50 to 6.34) that typifies the active state of β2AR^17^, and the positioning of residues R259 to L266 in ICL3 relative to the C-terminal end of TM7 (R328). In our prior research, we identified that this set of residues in ICL3 formed a turn and half of a helix and covered the G protein binding site as a loose “cap” (Fig. 1B). This cap motif has two distinct conformations: one that obstructs the G protein binding site, while the other one is open (shown in Fig. 1B, blue and green structure, respectively). The intracellular region of TM6 moves away from the intracellular region of TM3 upon activation of β2AR and this distance is used to assess the level activation of β2AR. Using these two metrics we classified the ten conformational clusters as follows:

- The active state, in which the ICL3 cap is extended out of the G protein binding site and the entire ICL3 is in an open state, and the TM6 moved away from TM3.
- The intermediate state, wherein TM6 remains away from TM3 but the ICL3 cap is loosely bound to the G protein coupling site.
- The inactive state. where TM6 is close to TM3 effectively occluding the G protein binding site. While the ICL3 cap is extended similarly to the active state, other portions of ICL3 stabilize the inactive state of the receptor through contacts with ICL2 and ICL1 residues. The fact that ICL3 contacts with ICL1/ICL2 in in active state and bound to the G protein binding site in the intermediate state make it a secondary regulator to G protein entry.

To sort the 10 clusters into these three main conformational states of β2AR described above, we used hierarchical clustering based on two parameters: the distance between residues 3.50 and 6.34, which reflects the TM3-TM6 distance, and the shortest distance between the ICL3 cap’s residues (R259 to L266) and R328 on the TM7 (Fig. 1C).The minimum distance between ICL3 cap residues and R328 distinctly differentiates intermediate states from active states with no overlap, leading us to select this as the metric to characterize the movement of the ICL3 cap. The hierarchical clustering identified three distinct clusters (Fig. 1D). The first cluster exhibits a small 3.50-6.34 distance and a medium ICL3-TM7 distance. Upon visualization, this conformation closely aligns with the fully inactive state EM structure^18^, leading us to label this cluster as the “inactive state ensemble”(Figs. 1B, 1C). The second cluster, with a large 3.50-6.34 distance and a small ICL3-TM7 distance, suggests that while the TM6 has shifted outwards, the ICL3 cap still obstructs the G protein binding site (Figs. 1B, 1C). We designate this as the “intermediate state ensemble”. The third cluster shows a large 3.50-6.34 distance and a large ICL3-TM7 distance, indicating that the TM6 has swung out and the ICL3 cap is out of the G protein coupling site, and this is the active state ensemble (Figs. 1B, 1C). Here after in the manuscript, we will refer to these three states.

### PIP2 competes with ICL1 and ICL2 to interact with ICL3 and maintain an equilibrium between the active and intermediate states

The multi-lipid bilayer model has an assortment of lipids that are unevenly dispersed between its upper and inner leaflets^6,7,16,19–22^. We calculated the radial distribution function of the different lipids from the center of mass (COM) of the receptor The simulation data indicate that the negatively charged lipid PIP2 displays higher concentrations close in proximity to β2AR (Fig. S2A) Notably, PIP2 is only present in the intracellular side. Furthermore, the number of PIP2 molecules close to the receptor is far higher in the active state compared to the inactive state (Fig. S2B). As PIP2 has been previously shown to influence membrane receptor activity^6,7,16,19–22^, we further investigated the influence of PIP2 on the conformation of ICL3.

Initially, we examined the role of PIP2 in stabilizing different conformations of ICL3. We identified the PIP2 molecule that form persistent contact (>20% simulation time) with residues on the IC loops, then we projected the COM of these PIP2 and the seven transmembrane helices on to the X-Y plane of the membrane as population density contour map. As shown in Fig. 2A, there is an increased density of PIP2 near the TM5/TM6/ICL3 region in both active and intermediate states. In the inactive states, PIP2 is found near the TM2/TM4/ICL1/ICL2 region. We calculated the interaction between PIP2 and ICL3, and the interactions of ICL3 with the other two IC loops, ICL1 and ICL2. In the inactive state of β2AR, there is a significant number of contacts between residues in ICL3 and ICL2 with more than 20% frequency (Fig. 2B), but no interaction between ICL3 and PIP2 (Fig. 2C). In the fully active state, ICL3 residues show persistent interactions with PIP2 (Fig. 2C) and decreased interactions with ICL2 and ICL1 (Fig. 2B). In the intermediate state ensemble, there is an increase in the number of persistent contacts between ICL3 with PIP2 (Fig. 2C) compared to the inactive state and concurrent decrease in interactions between ICL3 and both ICL1 and ICL2 (Fig. 2B). Thus, the interplay between of contacts of ICL1/ICL2 and PIP2 with ICL3 plays a pivotal role in stabilizing different conformational states of β2AR.

**Figure 2:**
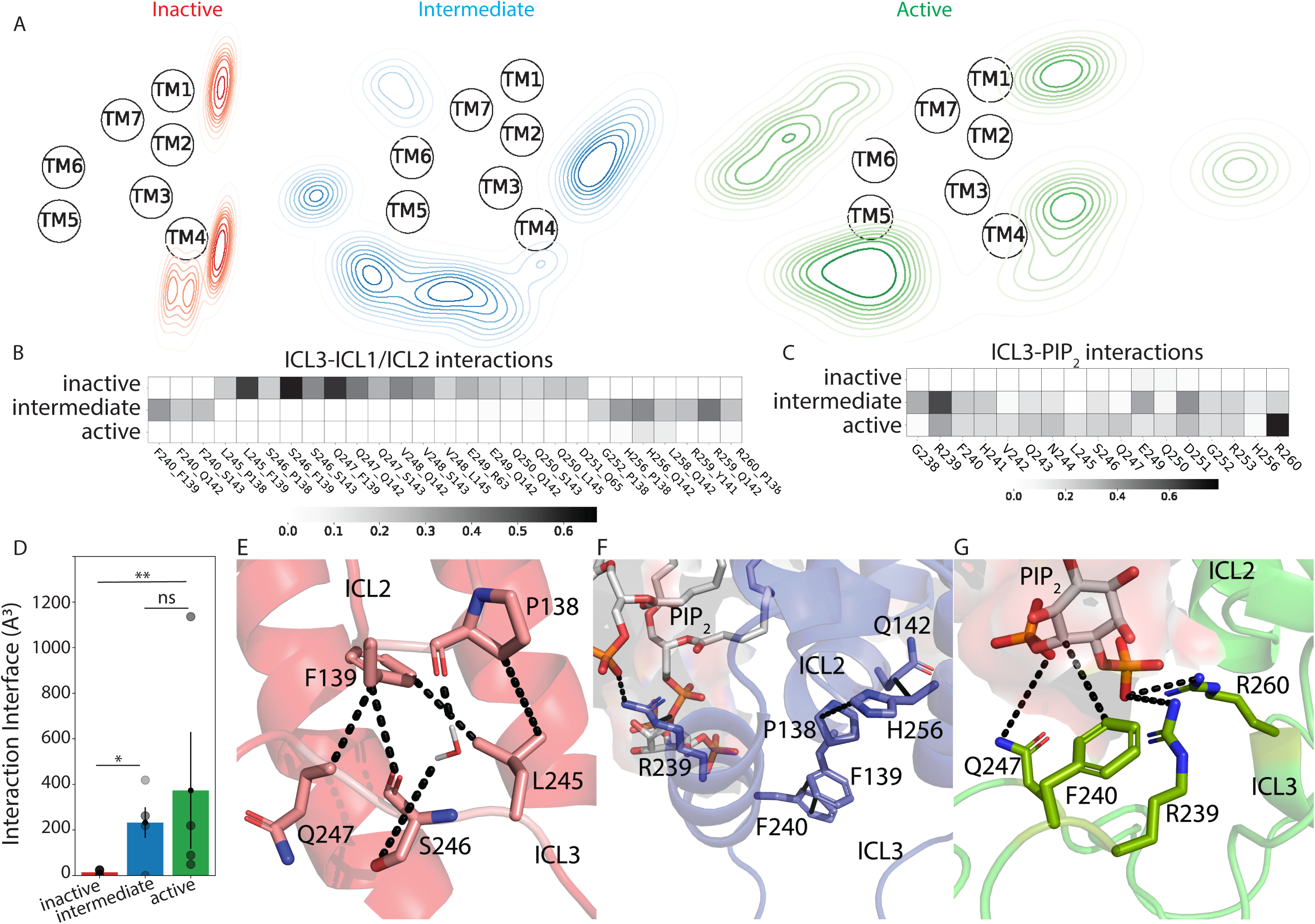
PIP2 competes with ICL1/2 for ICL3 binding and facilitate β2AR activation. A) Density of PIP2 distribution surrounding the seven-transmembrane domain for the three prime states. The contour visualization is derived by projecting the center of mass (COM) of the PIP2 head group onto the XY plane of the simulation box, with subsequent density calculations based on a defined grid. Red/Blue/Green contour map refers to the population density of PIP2 near the receptor in inactive/intermediate/active states. The COM of ICL1/ICL2 residues, and the center of mass of the middle of ICL3 (residue 250) is also projected on the XY plane. B) Frequency heatmap of the contacts between ICL3 and ICL1/2 residues. C) Frequency heatmap of the contacts between ICL3 residues and any PIP2 molecules. D) The interaction interface area between ICL3 and all the lipids that show contact persistence (>20%) with ICL3 residues. The ICL3-lipid interface area was calculated as the average value of all the 10 clusters for each of the conformational states. p-value for significance: ns – p=0.722, * - p=0.0278, ** - p=0.0079. E-G) The representative structure of ICL3 conformations in inactive/intermediate/active states respectively highlighting some of the residue contacts between ICL3-ICL1/2 or ICL3-PIP2.

We hypothesized that as ICL3 interacts favorably with PIP2, it will act as an anchor within the cell membrane, embedding the receptor deeper into the membrane bilayer. Supporting this, the contact surface area between ICL3 residues and all types of lipids is significantly higher in the active state than in the inactive state (Fig. 2D).

The exchange between interloop contacts and ICL3-PIP2 contacts accounts for this discrepancy (Fig. 2E-G). In the inactive state ensemble, F139 from ICL2 interacts with multiple residues L245, S246 and Q247 from ICL3 (Fig. 2E), forming a hydrophobic patch that covers the G protein binding site. There are also sporadic contacts (roughly 27.5% of the simulation time) between E249/D251 of ICL3 and R63/Q65 of ICL1. In the intermediate states, the contacts between L245, S246 and Q247 weaken. Instead, ICL3 residues F240, H256, R259 interact with ICL2 residues F139 and Q142. In the active state ensemble, there are no significant contacts between ICL3 and ICL1/2, the only less persistent pairs (< 20% simulation time) are between H256, L258 in ICL3 and Q142 in ICL2. While majority of the ICL3-ICL1/ICL2 contacts are hydrophobic in nature, the contacts between ICL3 and PIP2 are polar. In both active and intermediate states, the R239 and R260 form stable contacts with the negatively charged PIP2 head group (Fig. 2F-G). These ionic interactions anchor ICL3 in the membrane. Based on these results, we hypothesize that a competition exists between PIP2 and ICL1/2 for ICL3 interaction. When ICL3 binds with ICL1 and/or ICL2, it tends to favor more inactive states, whereas binding with PIP2 stabilizes active state ensemble.

### PIP2 anchoring ICL3 in the membrane bilayer tilts β2AR thus increasing the interaction interface of the receptor with the lipids

As described in the previous section, there is tight anchoring of ICL3 to PIP2 in the multi-lipid bilayer in the active state ensemble. This prompted us to study the effect of these anchoring interactions on the orientation of the receptor within the bilayer. To quantify this, we used the plane going across the middle of the lipid bilayer and computed the angle between the principal axis of β2AR and this plane (Fig. 3A). As shown in Fig. 3A, the distribution of the tilt angle for various states shows that both active and intermediate states have a pronounced tilt angle in comparison to the inactive state ensemble. This tilt causes the receptor to immerse deeper in the bilayer.

**Figure 3.**
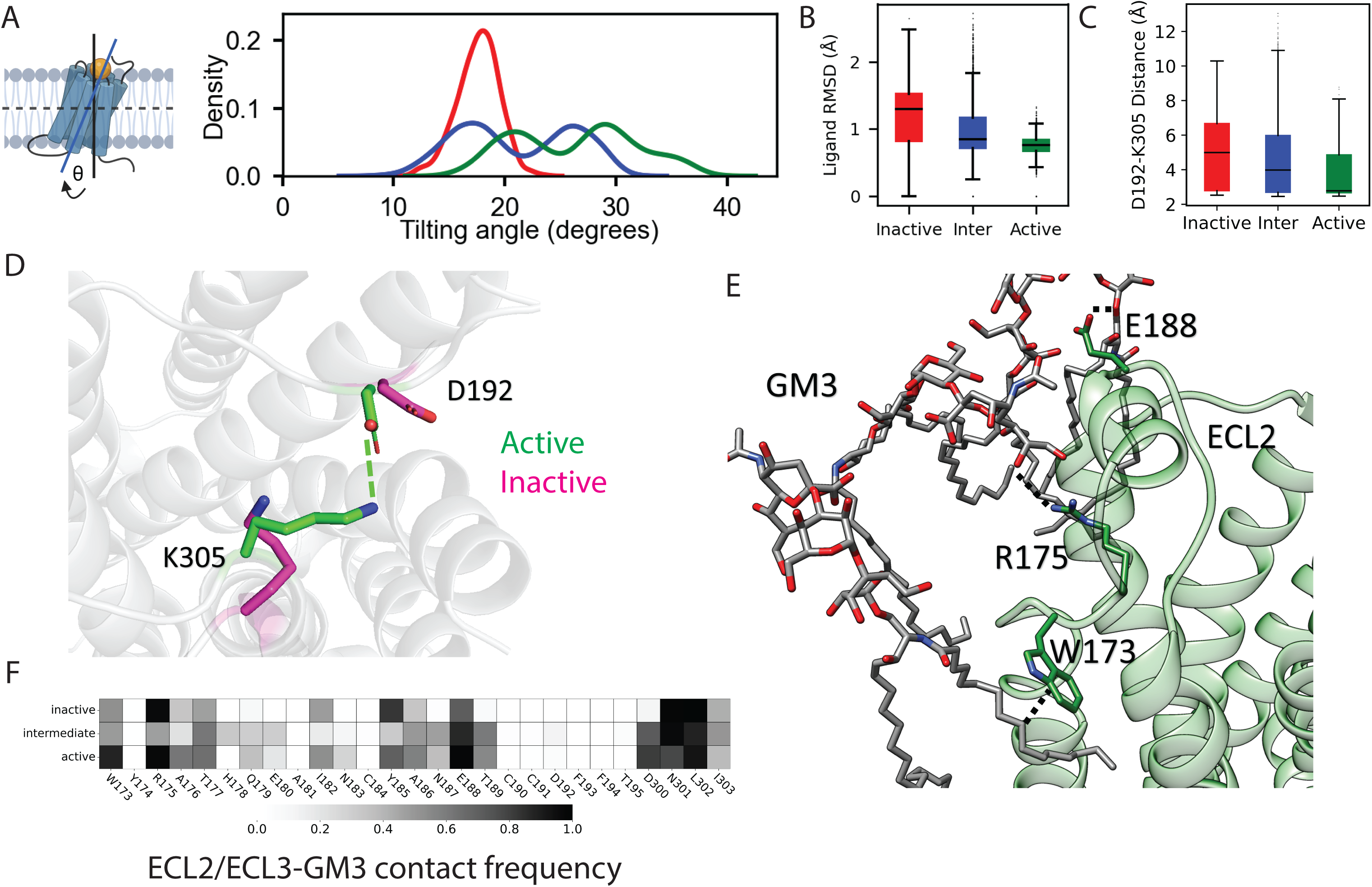
ICL3 anchoring in membrane lead to β2AR tilting, exerting an allosteric effect on the ligand binding pocket. (A) Tilting angle (θ) of GPCR in membrane for the inactive, intermediate and active states. (B) Root Mean Square Deviation (RMSD) of the coordinates of the agonist atoms in the inactive, intermediate, and active MD trajectories. (C) The minimum distance between D192 (ECL2) and K305 (ECL3) residue salt bridge in the inactive, intermediate, and active state MD trajectories. D) Three-dimensional structural representations highlighting the positions of D192 and K305. These amino acid residues are instrumental in maintaining the closed conformation between ECL2 and ECL3 regions. (E) Three-dimensional structural representation of the interaction between GM3 molecule and ECL2 residues. (F) Contact heatmap between GM3 and ECL2/ECL3 residues.

We studied the effect of the tilting motion seen in the active state of β2AR on the agonist binding site located in the extracellular region of β2AR. We analyzed the flexibility of the agonist, P0G (8-[(1R)-2-{[1,1-dimethyl-2-(2-methylphenyl)ethyl]amino}-1-hydroxyethyl]-5-hydroxy-2H-1,4-benzoxazin-3(4H)-one), in its binding site in the inactive, active and intermediate states. We calculated root mean square deviation (RMSD) in coordinates of the ligand over the entire MD ensemble as shown in Fig. 3B. The distribution of RMSD reveals that he agonist is less flexible in the active state ensemble (mean RMSD = 0.75 Å) compared to the inactive state ensemble (mean RMSD = 1.2 Å). Furthermore, we observed that the increased flexibility of the agonist in the inactive state comes from movement of the extracellular loops ECL2 and ECL3. Residues D192 in ECL2 and K305 in ECL3 form the salt bridge known to be important for ligand binding ^23^. The distance between these residues increases in the inactive state (5.1Å) compared to the active state (3.2Å), presumably weakening this salt bridge and increasing structural flexibility within the ECL2-ECL3 domain.

Our previous study demonstrated that the Ganglioside lipid (GM3) influences the dynamics of the extracellular loops of GPCRs^16^. This prompted us to explore the potential role of GM3 in facilitating the tilting of β2AR. A shown in Fig. S3 the density of GM3 projected on the XY plane of the membrane is high around the ECL2/ECL3 regions of β2AR in all the three states. Notably, the elongated sugar chain head of GM3 appears to flex, potentially covering the ECL2 region (Fig. 3E). By analyzing the contact frequency between residues in ECL2 and ECL3 and GM3, we identified that residues R175, E188, N301, and L302 show consistent contact with GM3 across all three states (Fig. 3F). However, a greater number of residues (12 residues) make persistent contacts (> 50% of the MD simulation snapshots) with GM3 in the active states compared to the inactive states (6 residues). Thus, GM3 could play an important role in holding the ligand in place in the receptor.

## Discussion

In this study we provide mechanistic evidence for the influence of lipid composition on the activity of β2AR predominantly through the ICL3 region of the receptor. This study supports previous findings concerning the influence of ICL3 on receptor activation through its broad conformational ensemble. We further delineated an important structural element of this ensemble, a short helical turn in the C-terminus of ICL3 that caps the G protein binding site. This “cap” forms when the receptor displays other structural hallmarks of activation (TM3-TM6 distance), indicating that ICL3 is a secondary regulator of the activation of β2AR. As shown in Fig. 4, we have shown that the negatively charged lipid, PIP2 interacts with the ICL3 residues in the active state of the receptor, thus opening up the ICL3 for G protein signaling. This finding is in agreement with previous findings that PIP2 is known to stabilize GPCR-G protein complex formation^6,7,16^. Sequence alignment of β2AR with GPCRs of similar ICL3 lengths reveals a degree of sequence similarity in both ICL2 and ICL3 regions (see Fig. S4). This similarity suggests that the competitive interaction observed between ICL2 and PIP2 for ICL3 may also be present in other GPCRs.

**Figure 4.**
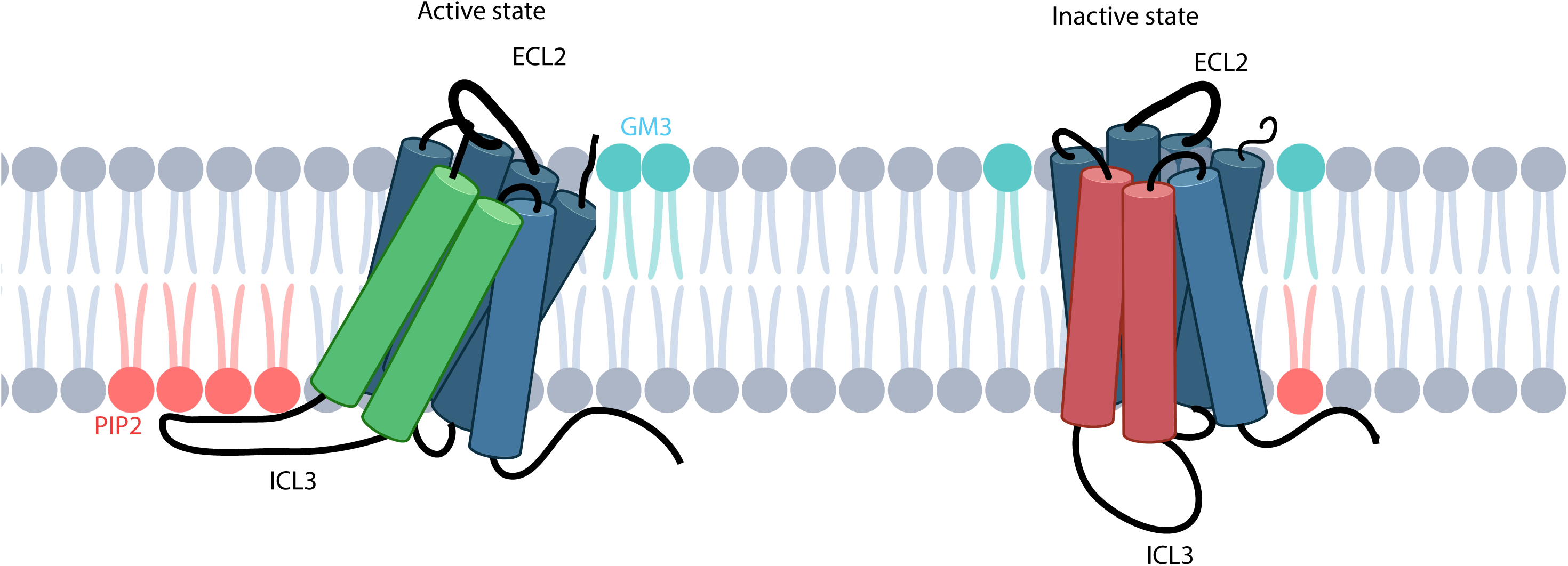
PIP2 anchors the active state of β2AR in the bilayer leading to stabilization of the open state of ICL3. This results in tilting of the receptor. GM3 on the outer leaflet of the membrane interacts strongly with ECL2 in membrane facilitating β2AR tilting, which further constrains the ligand binding site.

This PIP2-ICL3 anchor concerts an allosteric effect on the receptor through tilting of the receptor in the plasma membrane, stabilization of extracellular loop conformation, and ultimately stabilization of agonist binding. This stabilization is further coordinated by interactions between ECL2 and GM3, a lipid which is predominantly in the upper leaflet of the membrane^6,7,16^ and is known to regulate GPCR conformational dynamics^16,24^. Our latest findings not only reinforce the assertion that lipids are integral to GPCR function but also offer new perspectives on the mechanisms through which these lipids regulate receptor activity. In summary, our study provides deep mechanistic insight into role of lipids and therefore the role of cell membrane components on the regulation of receptor activity by intrinsically disordered loop regions in GPCRs.

## Methods

### MD simulations trajectories preparation

In our previous study^14^, we performed 22 μs of MD simulations to sampling the conformational heterogeneity of the full length ICL3 in β2AR. The absence of ICL3 in any resolved crystal or cryo-EM structures necessitated the use of computational modeling to construct the ICL3 loop. We generated four models of β2AR incorporating ICL3 (Fig. S5 and Table S1). Model A was a homology model of β2AR derived using the turkey β1AR inactive state structure (PDB ID: 2YCX), including the ICL3 loop. For Model B, the intermediate state crystal structure (PDB ID: 6E67)^25^ served as the template, with the entire ICL3 loop modeled to mirror the template structure. In Model C, while also using the intermediate state structure (PDB ID: 6E67)^25^ as the foundation, we constructed the C-terminal portion of ICL3 as an alpha-helix and the remaining segment as a random coil. Model D, based on the same intermediate structure (PDB ID: 6E67)^25^, features the ICL3 cap as an alpha-helix, with the rest of the loop modeled as a random coil. Details of the MD simulations are in reference 14.

To streamline the trajectory data for subsequent analysis, our strategy is two-fold. Firstly, we seek to condense the expansive 22-millisecond trajectory, which is currently too extensive and may introduce unnecessary variability into our findings. To mitigate this and assure the integrity of our results, we implemented principal component analysis (PCA) on the root-mean-square deviation (RMSD) of the GPCR backbone. This technique allowed us to organize the trajectory into a number of distinct clusters. From each cluster, we then selected a small subset of conformations situated near the population density wells to act as representative states for the entire simulation. Our second aim is to draw a connection between these representative states, derived from the clustering process, and specific states that define GPCR activity. This will enable us to elucidate the influence of varying ICL3 conformations on the regulation of GPCR activity. The method employed in both steps will be elaborated in the following section.

### Principal component analysis on GPCR backbone and acquire sub cluster trajectories

We utilized the 22 μs simulation trajectory from our prior research^14^ and conducted a PCA based on their backbone RMSD. Subsequently, we projected the first two principal components (PC1 and PC2) onto an X-Y plane and established the population density contour map by using Seaborn KDE plot (Figure 1A)^26^. To effectively group the sample points into distinct sub-states, we employed Kmeans clustering based on the PC1 and PC2 values. We iteratively set n_clusters from 2 to 20 and computed both the Silhouette score and the Davies-Bouldin index for each iteration. The results, as depicted in Figure S1A and S1B, indicate that a n_clusters value of 12 yields a high Silhouette score and a low Davies-Bouldin index. Consequently, we determined that n_clusters=12 is the optimal parameter for this clustering analysis.

After determining the 12 clusters, we assessed the population density by categorizing all data points into a 35×35 bins grid based on their PC1 and PC2 values. Within each of the 12 clusters, we identified the bin with the greatest population density. Only the frames from these high-density bins were selected for subsequent analysis. Among the 12 identified clusters (Figure S1C), clusters 2 and 4 have sparse sampling points and do not display any global population density maximum as illustrated in Figure 1A. Consequently, we chose to exclude these two clusters and directed our attention solely to the remaining 10 clusters. From each of the 10 clusters, we selected the frames situated at the centers of their respective bins. These frames serve as representative structures and the centroid structures of each cluster are displayed as three-dimensional configurations in Figure 1A.

### Distance measurement and hierarchical clustering

The distance measurements were conducted using the MDAnalaysis distance module^27^. Specifically, the distance between residues 3.50-6.34 was determined between the Ca atoms of residue 131 and 272, in line with the BW numbering table provided by GPCRdb.org. The TM7-ICL3 cap distance was gauged as the minimum distance from R328 on TM7, to the center of mass (COM) of residues 259 to 266 in ICL3. Hierarchical clustering was executed on the distances of 3.50-6.34 and the TM7-ICL3 cap from the 10 clusters, employing the single linkage method available in the scipy.cluster.hierarchy module. The D192-K305 distance was measured as the minimum distance between residue 192 and residue 305.

### Contact analysis

In this study, all contact analyses were carried out using get_contact <u>(get_contact.io</u>). For Figure 2B, residues from ICL1, ICL2, and ICL3 were selected based on the BW table on GPCRdb.com. For Figure 2C, ICL3 residues and all PIP2 molecules present in the MD simulation were chosen. Meanwhile, for Figure 3I, residues from ECL2 and ECL3 as well as all GM3 molecules in the MD simulation were selected. The contact calculations were executed using the default parameters of get_contact. All types of contacts, such as van der Waals contacts, hydrogen bond contacts, and π-π contacts, were factored into the calculations.

### Projecting lipid/GPCR onto X-Y plane

To generate lipid distribution contour maps as depicted in Figure 2A and 3G, several procedures were executed. Firstly, the protein structures were aligned using their Ca atoms. Next, the COM for both the seven TM regions and the head groups of the targeted lipids, PIP2 and GM3, was determined. The TM region range was defined based on the BW numbering table from GPCRdb.com. To define the orientation of the GPCR, the average X and Y coordinates of the seven TMs COM were projected onto the membrane X-Y plane. Additionally, the X and Y coordinates of each lipid molecule that consistently interacted (>20% simulation time) with either the intracellular loops (ICLs) or extracellular loops (ECLs) were also projected onto this plane. Through these processes, the spatial distribution and orientation of lipids relative to the GPCR became visible, providing a detailed insight into their interactions and placement.

### Interaction interface area calculation

We employed the FreeSASA python module (www.freesasa.github.io), to compute the interaction interface area. Initially, we determined the total solvent-accessible surface area (SASA) for either ICL3 or ECL2 and all lipids within the MD simulation. Subsequently, the SASA was calculated separately for the ICL3 or ECL2 region and for all lipids. The interaction interface area is deduced by taking the sum of SASA for ICL3 or ECL2 and SASA for the lipids, and then subtracting the total SASA from this sum. We then computed the average interaction interface area for each of the 10 clusters independently. In Figure 2D and 3J, each cluster is represented by a single dot, indicating its average interaction interface area. A Mann-Whiteney U test (scipy.states.mannwhitneyu) was performed on the interaction interface area values to test the significancy. Given that the inactive state only has one cluster, we extracted average interaction interface area values from five distinct segments of inactive cluster, each consisting of consecutive frames of 20% of the total simulation length. This segmentation approach provides us sufficient data points for Mann-Whitney U test.

### Membrane plane fitting measurement

To calculate the tilting angle, we utilized a comprehensive approach, fitting a plane through all lipid molecules in the membrane. Principal Component Analysis (PCA) was employed to facilitate this fitting, with the smallest principal component defining the plane’s normal vector. Subsequently, the principal axis of the protein was determined by fitting it to the backbone of the transmembrane helices, enabling the calculation of the tilting angle—the angle between the plane’s normal vector and the protein’s principal axis.

### Radial pair distribution function G(r) calculation

The radial pair distribution function, denoted as G(r), was determined with the help of the VMD G(r) module^2827^. Using the center of mass (COM) of the entire GPCR as the central reference, we adopted a bin size of 0.5 Å for the radial pair. The calculation for G(r) spanned from 0 to 30 Å, and this was executed for all ten lipid types.

### Ligand RMSD calculation

The ligand RMSD was calculated by using MDAnalysis.rms module^27^. We first align the protein by Ca atoms of TM region, using the initial model as reference. Then we calculate the RMSD of ligand per frame. All RMSD values of each state, after combining the corresponding clusters, were plotted in bar graph form in Figure 3C.

### Declaration of generative AI and AI-assisted technologies in the writing process

During the preparation of this work the author(s) used ChatGPT-4.0 in order to revise the language. After using this tool/service, the author(s) reviewed and edited the content as needed and take(s) full responsibility for the content of the publication.

## Acknowledgements

This research was funded by the NIH (R35-GM126940 to S.S., R01-GM117923 to N.V. and T32-AR007612 to F.S.).

