## Supplemental Information for "Cellular Lipids Regulate the Conformational Ensembles of the Disordered Intracellular Loop 3 in β2 Adrenergic Receptor"

**Supporting Information**

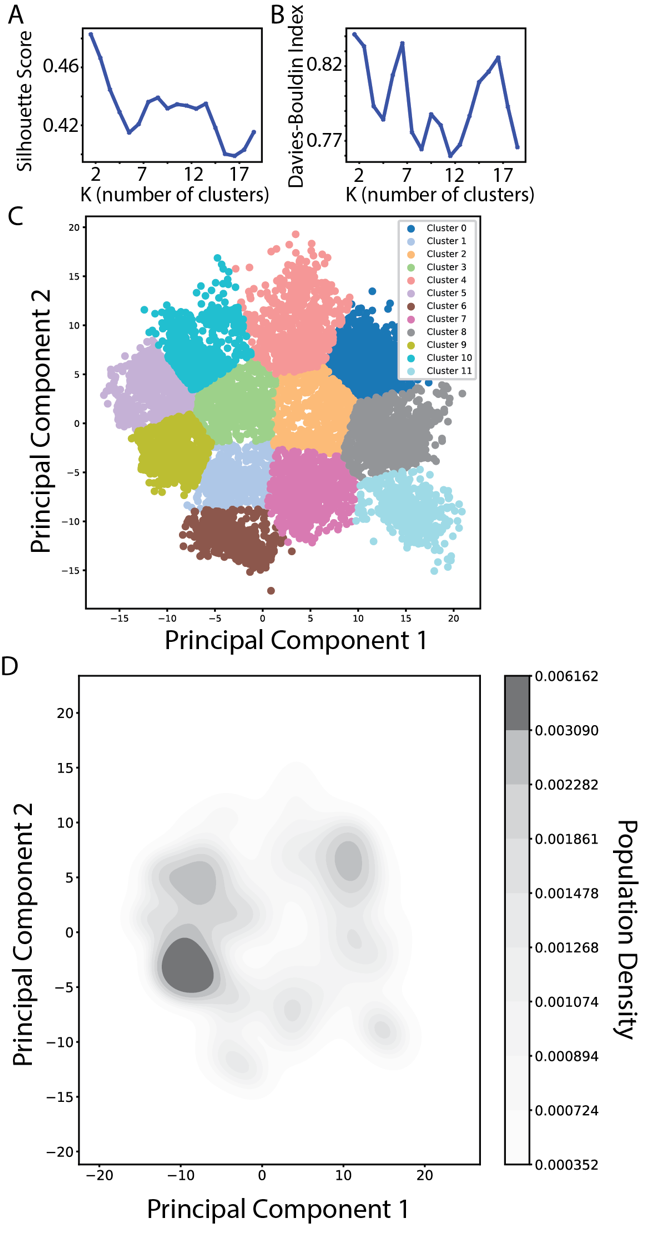

**Figure S1**. A, B) Silhouette scores and Davies-Bouldin index plots depict the optimal number of clusters. An elevated Silhouette score combined with a reduced Davies-Bouldin index indicates more effective clustering. When setting n_clusters to 12, both the Silhouette score in panel A and the Davies-Bouldin index in panel B present the most optimal combination, suggesting that n_clusters=12 is the ideal parameter choice for clustering. C) A scatter plot representation of the clustering outcome using n_clusters=12. Among these 12 clusters, cluster 4 and cluster 2 lack population density maxima and were omitted from further analysis. D) The population density landscape from Figure 1A for comparison.

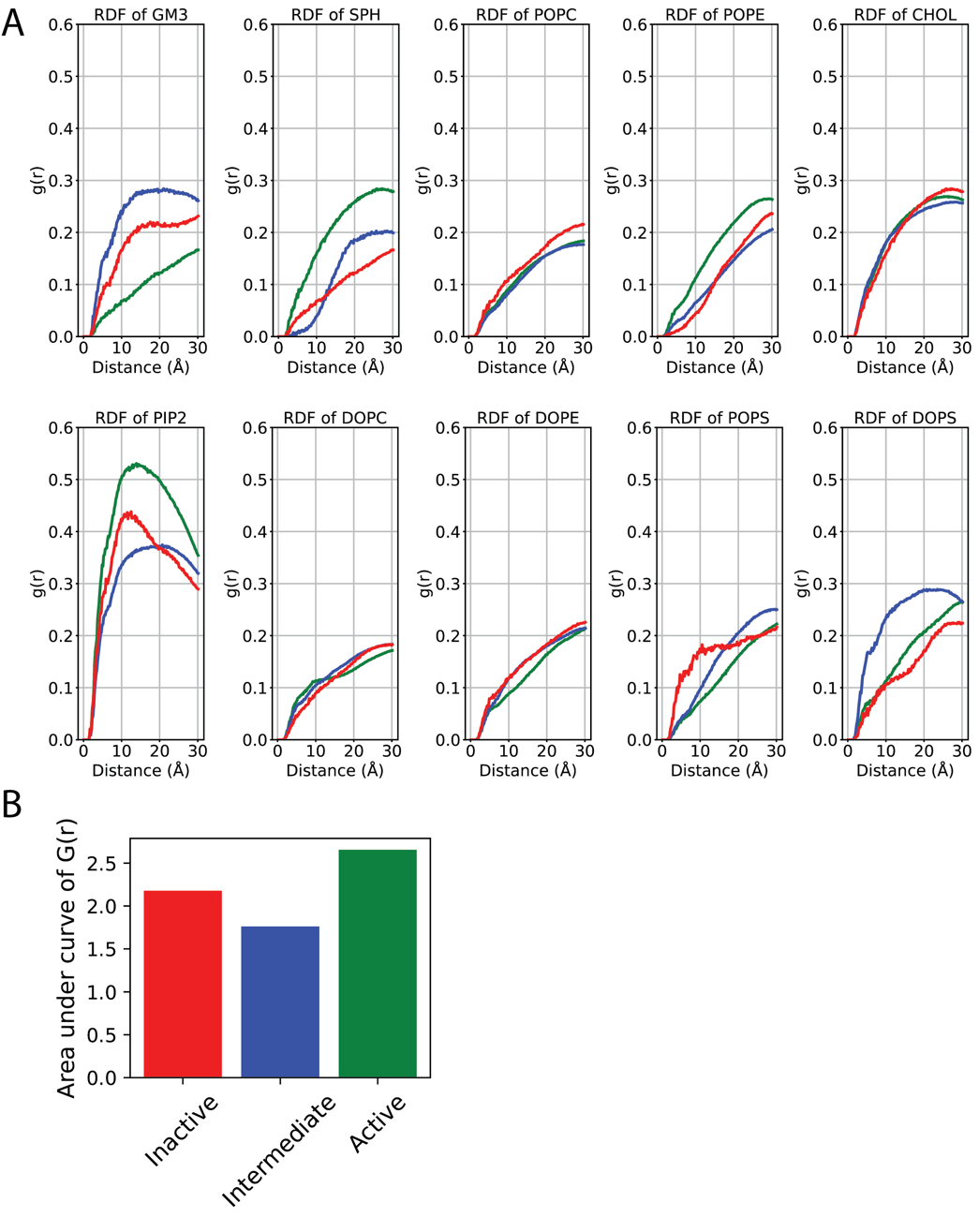

**Figure S2**: A. Radial distribution function (G(r)) for different lipids in the three major receptor states. Red curve is for inactive state, blue curve is for intermediate state, green curve is for active state. PIP2 shows relatively high G(r) at 10 Å distance to the center of mass of β2AR. B. The area under the G(r) curves within 10 Å of the receptor center of mass for PIP2 in the inactive, intermediate, and active states.

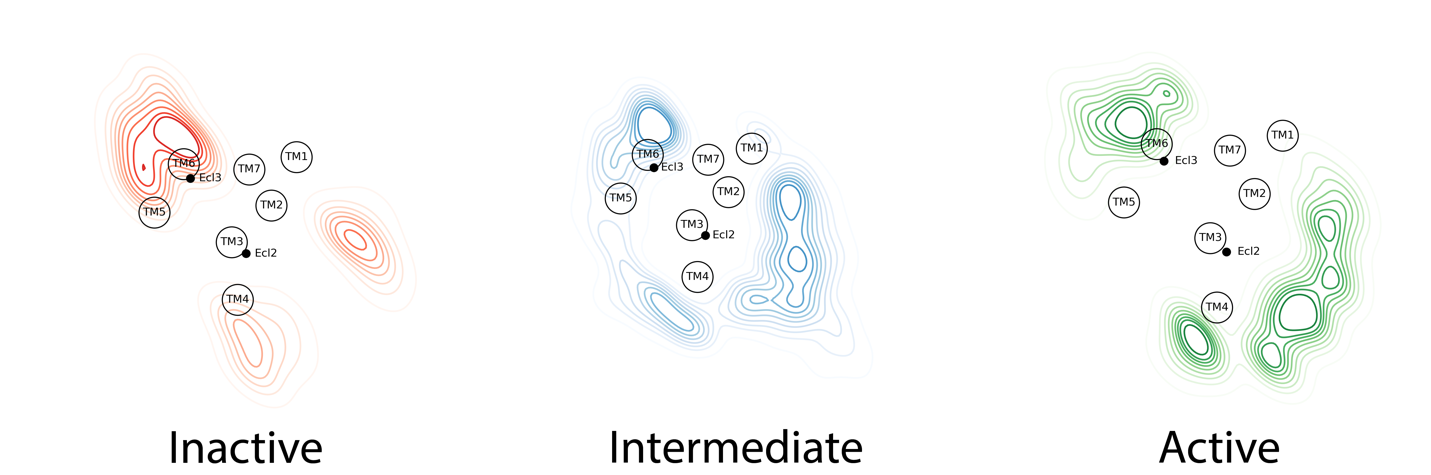

**Figure S3**. The density of GM3 around each transmembrane region and extracellular loops in the outer leaflet of the multi-lipid bilayer in the three conformational states of β2AR. The contour visualization is derived by projecting the COM of GM3 head group atoms onto the XY plane of the simulation box, with subsequent density calculation based on a defined grid.

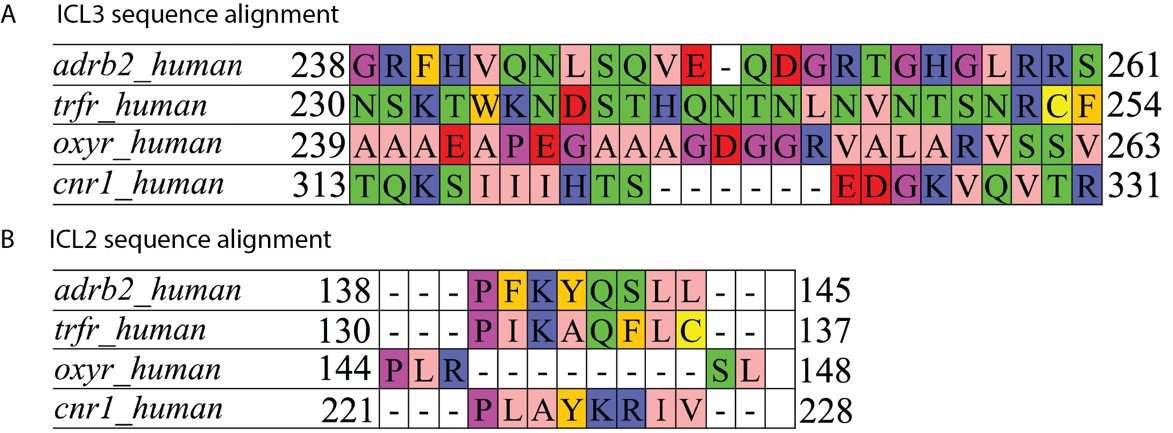

**Figure S4**. Sequence alignment of ICL2 and ICL3 between β2AR and GPCR with similar length of ICL3.

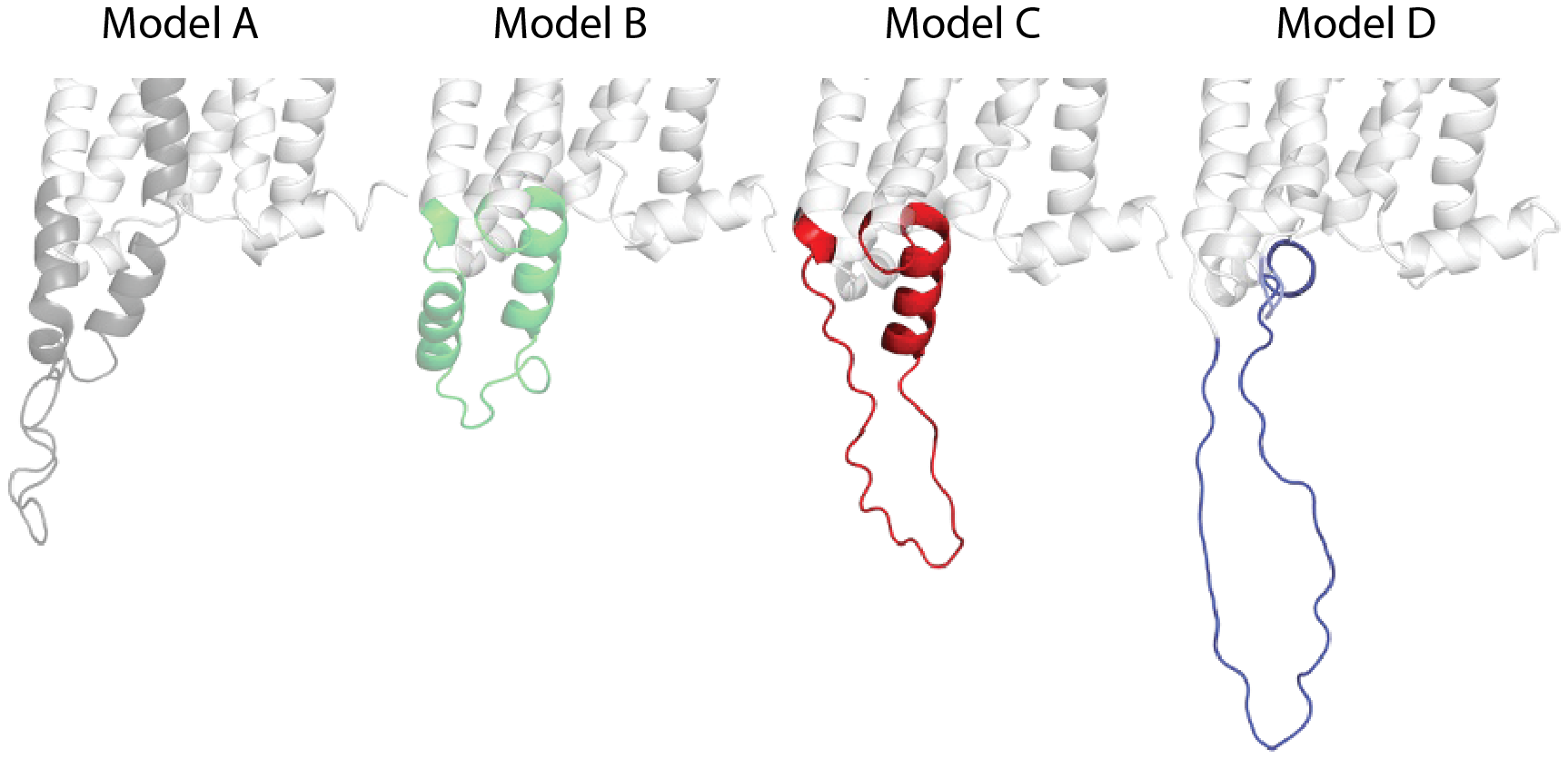

**Figure S5**. Structural models of ICL3 in β_2_AR used as starting structures for MD simulations in cell membrane conditions.

**Supplementary Table 1**. Simulations from previous study. Refer to previous study for model description (14).

| Simulations performed | **Random velocity used** | **Simulation time for each velocity (μs)** | **Total simulation time  (μs)** |
| --- | --- | --- | --- |
| Model A | 5 | 0.4 | 2 |
| Model B | 5 | 0.4 | 2 |
| Model C | 5 | 0.4 | 2 |
| Model D | 25 | 0.4 | 10 |
| Extension simulation from Model D, 4^th^ velocity to fill the gap on population density landscape. | | | |
| Starting structure: snapshot at end of 50 ns of Model D 4^th^ velocity | 5 | 0.4 | 2 |
| Starting structure: snapshot at end of 100 ns of Model D 4^th^ velocity | 5 | 0.4 | 2 |
| Starting structure: snapshot at end of 150 ns of of Model D 4^th^ velocity | 5 | 0.4 | 2 |
|  |  |  | Sum = 22 |

**Supplementary Table 2**. The lipids composition in the mixed lipid bilayer.

|  | POPC | DOPC | POPE | DOPE | POPS | DOPS | GM3 | Sph | PIP2 | CHOL |
| --- | --- | --- | --- | --- | --- | --- | --- | --- | --- | --- |
| Upper | 20% | 20% | 5% | 5% | 0% | 0% | 10% | 15% | 0% | 25% |
| Lower | 5% | 5% | 20% | 20% | 7% | 8% | 0% | 0% | 10% | 25% |
